# Elevated risk of invasive group A streptococcal disease and host genetic variation in the human leukocyte antigen locus

**DOI:** 10.1101/559161

**Authors:** Tom Parks, Katherine Elliott, Theresa Lamagni, Kathryn Auckland, Alexander J. Mentzer, Rebecca Guy, Doreen Cartledge, Lenka Strakova, Daniel O’Connor, Andrew J. Pollard, Matthew J. Neville, Anubha Mahajan, Houman Ashrafian, Stephen J. Chapman, Adrian V. S. Hill, Shiranee Sriskandan, Julian C. Knight

## Abstract

Invasive group A streptococcal (GAS) disease is uncommon but carries a high case-fatality rate relative to other infectious diseases. Given the ubiquity of mild GAS infections, it remains unclear why healthy individuals will occasionally develop life-threatening infections, raising the possibility of host genetic predisposition. Here, we present the results of a case-control study including 43 invasive GAS cases and 1,540 controls. Using HLA imputation and linear mixed-models, we find each copy of the *HLA-DQA1*\*01:03 allele associates with a two-fold increased risk of disease (odds ratio 2.3, 95% confidence interval 1.3-4.4, *P*=0.009), an association which persists with classical HLA typing of a subset of cases and analysis with an alternative large control dataset with validated HLA data. Moreover, we propose the association is driven by the allele itself rather than the background haplotype. Overall this finding provides impetus for further investigation of the immunogenetic basis of this devastating bacterial disease.

## Introduction

Invasive group A streptococcal (GAS) disease is defined by isolation of *Streptococcus pyogenes* at a normally sterile site. Although uncommon, the incidence rate reaching 3 per 100,000 in Northern Europe^1^, the case-fatality rate is high relative to other infections, reaching 20% in some studies.^2^ While infection can occur at a variety of sites, soft tissue infections predominate, of which necrotising fasciitis (NF) is a rare but particularly dangerous form often necessitating extensive surgical debridement. This and other forms of invasive GAS disease can be complicated by streptococcal toxic shock syndrome (STSS) characterised by hypotension, multi-organ failure and a case-fatality rate exceeding 40%^1^.

Despite growing recognition of the importance of host genetic factors in susceptibility to infectious diseases, limited attention has so far been paid to host genetic susceptibility to invasive GAS disease.^3^ The only study to investigate this in humans dates from the candidate gene era focussing on haplotypes in the class II region of the human leukocyte antigen (HLA) locus^4^. Rather than investigating susceptibility itself, this study reported specific haplotypes associated with severe disease defined by the presence or absence of hypotension and multi-organ failure. In particular, the authors found the *HLA-DRB1*\*1501/*HLA-DQB1*\*0602 haplotype to be associated with a four-fold reduced risk of severe disease among previously healthy individuals with invasive GAS disease.^4^ Nonetheless, further support for the role of HLA in invasive GAS comes from several studies showing binding of GAS superantigens to HLA-DQ molecules.^5^ In particular, streptococcal pyrogenic exotoxin A (SpeA), a key superantigen, binds HLA-DQA1 in a manner dependent on DQA1 polymorphism.^6^ Added to this, transgenic mice expressing human HLA-DQ molecules were found to be highly sensitive to superantigens compared to non-transgenic littermates^7^, while particular HLA-DQ molecules were associated with enhanced infection of the nasal cavity in a manner dependent on SpeA.^8^ Nonetheless, despite rapid recent progress in the field of human genetics, the association between the HLA locus and invasive GAS has not been revisited, likely reflecting the challenges of recruiting patients with what is essentially a rare and extreme phenotype.

In the present study, we investigate the relationship between HLA class II alleles and susceptibility to invasive GAS disease, limiting our analysis to otherwise previously healthy children and young adults. Here, using contemporary methods that are robust to the major confounders of candidate gene approaches, we find the *HLA- DQA1*\*01:03 allele to be associated with a two-fold increased risk of susceptibility to invasive GAS disease. While this allele is not part of any of the haplotypes linked to the trait in the candidate gene era^4^, it adds weight to the notion that HLA polymorphism contributes to the outcome of invasive GAS infections, perhaps in a manner dependent on GAS superantigens.^6^ Overall this finding provides impetus for further investigation of the immunogenetic basis of this devastating bacterial disease.

## Results and Discussion

After quality control, we included 43 cases of European ancestry aged less than 65 years without comorbidity (Supplementary Table 1). Of these, 34 had been diagnosed with NF while nine had been diagnosed with other manifestations of invasive GAS disease (Table 1). The youngest patient was 18 months and the eldest was 63 years (median 35 years, interquartile range 25-41 years). Four of the seven children had preceding varicella, five of the women were postpartum, two after caesarean section, and two other adult patients after other surgery. Otherwise, the patients had no risk factors for invasive GAS disease. For our primary case-control analysis we compared our cases with 1,540 healthy children of European ancestry previously recruited to studies of vaccine efficacy undertaken by the Oxford Vaccine Group, University of Oxford, Oxford, UK. For sensitivity analyses, we compared our cases with 430 healthy adults of European ancestry, a subset of the 5,544 individuals from the National Institute for Health Research Oxford Biobank, for whom validated HLA data were available.^9^

**Table 1.**
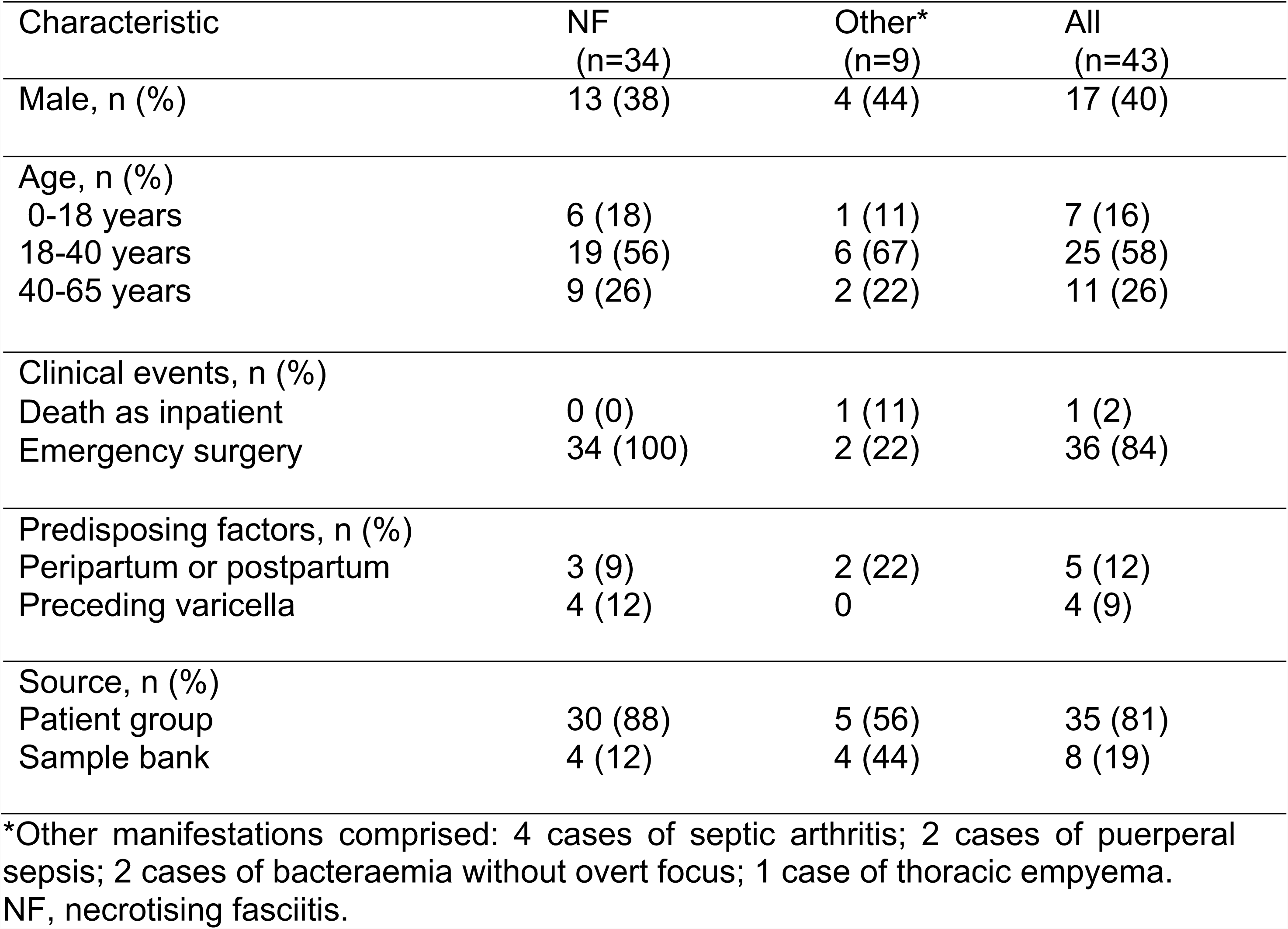
Clinical characteristics of invasive GAS cases

| Characteristic | NF<br>(n=34) | Other*<br>(n=9) | All<br>(n=43) |
| --- | --- | --- | --- |
| Male, n (%) | 13 (38) | 4 (44) | 17 (40) |
| Age, n (%) |  |  |  |
| 0-18 years | 6 (18) | 1 (11) | 7 (16) |
| 18-40 years | 19 (56) | 6 (67) | 25 (58) |
| 40-65 years | 9 (26) | 2 (22) | 11 (26) |
| Clinical events, n (%) |  |  |  |
| Death as inpatient | 0 (0) | 1 (11) | 1 (2) |
| Emergency surgery | 34 (100) | 2 (22) | 36 (84) |
| Predisposing factors, n (%) |  |  |  |
| Peripartum or postpartum | 3 (9) | 2 (22) | 5 (12) |
| Preceding varicella | 4 (12) | 0 | 4 (9) |
| Source, n (%) |  |  |  |
| Patient group | 30 (88) | 5 (56) | 35 (81) |
| Sample bank | 4 (12) | 4 (44) | 8 (19) |

\*Other manifestations comprised: 4 cases of septic arthritis; 2 cases of puerperal sepsis; 2 cases of bacteraemia without overt focus; 1 case of thoracic empyema. NF, necrotising fasciitis.

We first considered genotypic associations in the extended major histocompatibility complex based on SNP genotyping. Among 434 directly ascertained genotypes with minor allele frequency (MAF) greater than 5%, the strongest association signal was found at rs2534816 (*P*_LMM_=0.0013) located in the class I region 37kb from *HLA-E*. The strongest signal among 137 variants in the class II region was found at rs9276171 (*P*_LMM_=0.006) located in the intron of *HLA-DQB3*. Of 4,585 imputed genotypes the strongest signal was found at rs2524222 (*P*_LMM_=0.0005), located 49kb from *HLA-E*, while the strongest class II signal was found at rs1383265 (*P*_LMM_=0.004), located 8.5kb from *HLA-DQB2* (Supplementary Figure 1).

We then proceeded to analyse associations based on HLA imputation. Of 160 imputed four-digit HLA alleles, the strongest signal was linked to *HLA-DQA1*\*01:03 allele (Figure 1A), which was found at MAF 12.7% in cases compared to 5.9% in controls (odds ratio, OR, 2.3, 95% confidence interval, CI, 1.2-4.4, *P*_LMM_=0.009). Consistent with this, the presence of a lysine in place of an arginine at position 41, corresponding to rs36219699, and an alanine in place of a serine at position 130, corresponding to rs41547417, which together define *DQA1*\*01:03, were similarly associated with disease (*P*_LMM_=0.009). The *DQA1*\*01:03 signal was marginally weaker when limiting the analysis to the 34 patients with NF (OR=2.1, 95% CI 1.0-4.4, *P*_LMM_=0.049) despite the fact it was preserved in an analysis based on the similarly sized subgroup of 32 patients aged less than 40 years (OR=2.6, 95% CI 1.3-5.2, *P*_LMM_=0.007). Two additional four-digit alleles and three additional amino acids in the class II region, along with five amino acids in the class I region were associated with susceptibility with *P*_LMM_ less than 0.05 (Supplementary Tables 2 & 3). However, after controlling for the presence of *DQA1*\*01:03, none of the four-digit class II alleles remained significant at this level (Figure 1B).

**Figure 1.**
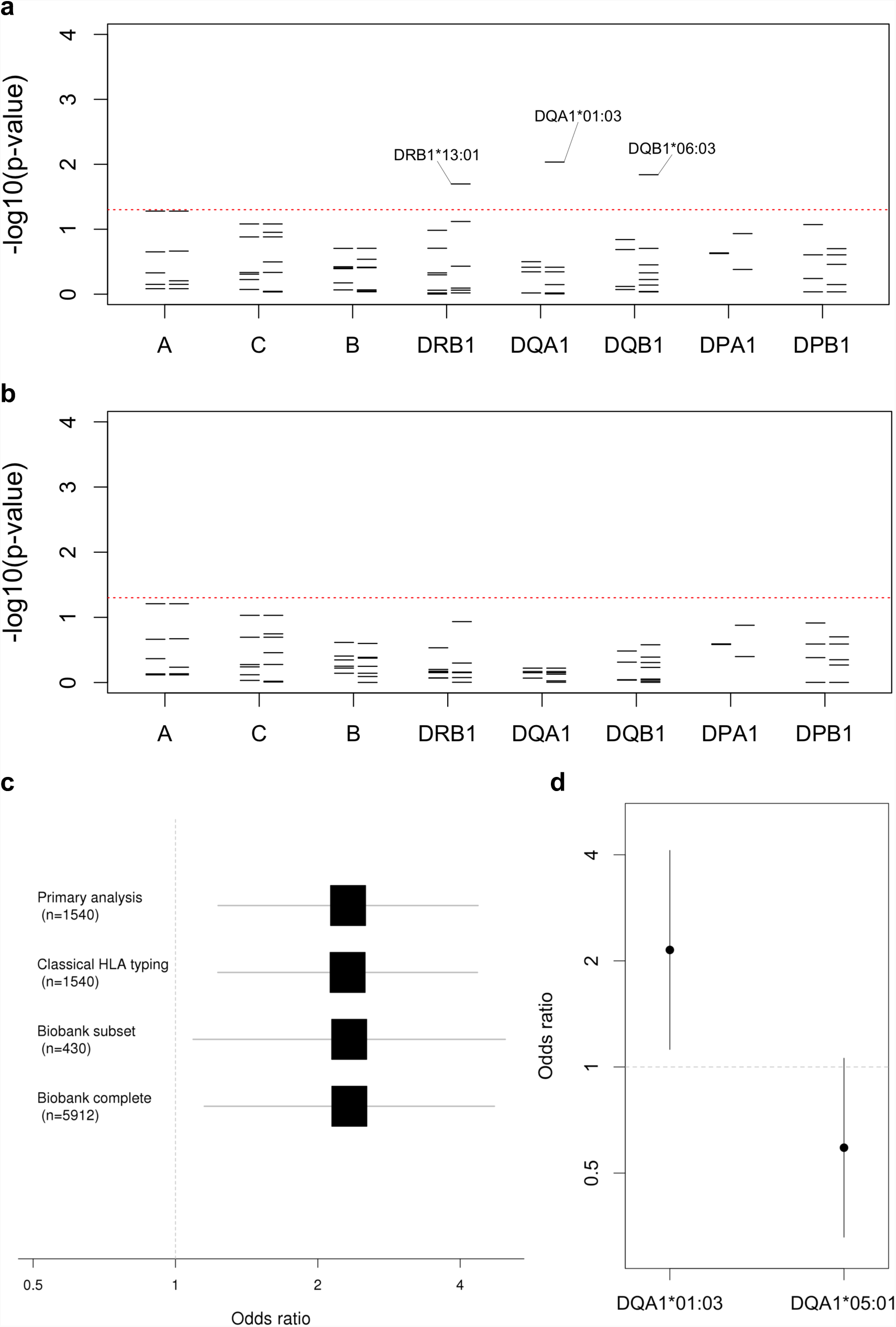
Classical HLA alleles associated with invasive GAS disease. **(a)** For each locus the negative common logarithm of the p-value from LMM analysis is plotted with two-digit alleles to the left and four-digit alleles to the right. **(b)** The same is plotted for an LMM analysis conditioned on *HLA-DQA1*\*01:03. **(c)** For the primary analysis and three sensitivity analyses, effect size estimates for *HLA-DQA1*\*01:03 are shown based on a logarithmic scale with the number of controls in each shown in brackets. The first three analyses use LMM with transformation^36^, while the latter based on the entire Oxford Biobank uses logistic regression. **(d)** For both *HLA-DQA1*\*01:03 and *HLA-DQA1*\*05:01, effect size estimates are shown, on a logarithmic scale, each conditioned on the other allele.

To validate our findings, we performed classical HLA typing of the *DRB1, DQA1* and *DQB1* loci in 30 cases for which sufficient DNA was available. Across the 42 alleles observed, concordance with imputation was generally high, ranging from 85.0% for *DRB1* through 91.7% for *DQA1* to 96.7% for *DQB1*. Moreover, the six copies of *DQA1*\*01:03 were perfectly imputed while only three of the remaining alleles were imputed with accuracy of 95% or less. We then reran our analyses substituting the available classical for imputed HLA types and found the effect size estimate for *DQA1*\*01:03 unchanged (Figure 1C).

We next repeated our analyses comparing the 43 cases to the alternative population of adult European controls from the Oxford Biobank among whom the MAF of *DQA1*\*01:03 was also 5.9%. The effect size estimate for *DQA1*\*01:03 remained unchanged in analyses using either logistic regression with all 5,544 European individuals for whom validated HLA data were available, or a linear mixed model with the subset of 430 European individuals for whom genome-wide data were available (Figure 1C), the latter correcting for ancestry and relatedness.

Next, we investigated effects of other *DQA1* alleles using both linear mixed-models and logistic regression. Based on likelihood ratio, the best fit was achieved by a model comprising fixed effects parameters for both *DQA1*\*01:03 and *DQA1*\*05:01 (Figure 1D), the latter having a weak protective effect (OR=0.62, 95% CI 0.35-1.09). In this scenario, each copy of *DQA1*\*01:03 was associated with a two-fold increased risk of invasive GAS disease (OR=2.1, 95% CI 1.2-4.1), an effect size and allele frequency that would imply a population attributable fraction of 11.6%. Additionally, the effect size estimates for the two alleles were highly consistent across a number of alternative analytical approaches including logistic regression with or without principal components^10^ and a generalised linear mixed-model analysis (Supplementary Figure 2), also termed logistic mixed-model analysis.^11^

Finally, we investigated whether the signal was better explained by haplotypes or individual alleles. Having defined nine three-locus class II haplotypes with MAF greater than 5%, we tested their association with susceptibility. Of these only the *DRB1*\*13:01-*DQA1*\*01:03-*DQB1*\*06:03 haplotype, which had a MAF of 11.6% in cases compared to 5.6% in controls, was significantly associated with susceptibility (OR 2.2, 95% CI 1.2-4.3, *P*_LMM_=0.015). Interestingly, one copy of the rarer *DRB1*\*15:02-*DQA1*\*01:03-*DQB1*\*06:01 was also present among the cases giving a MAF 1.1% compared to 0.23% among controls, although this difference was not statistically significant (OR=5.1, 95% CI 0.8-34, *P*_LMM_=0.09). We did not observe the *DQA1*\*01:03 allele in any other haplotype with MAF down to 0.01%. No further class II alleles with MAF greater than 5% were associated with susceptibility, including the previously implicated *DRB1*\*15:01-*DQA1*\*01:02-*DQB1*\*06:02 haplotype^4^, which was present in 14.0% of cases and 15.8% of controls (*P*_LMM_=0.67). However, consistent with the same earlier report, the *DRB1\**14:01-*DQA1*\*01:01-*DQB1\**05:03 haplotype, with MAF 5.8% in cases compared to 2.5% in controls, was associated with increased risk of disease (OR=2.4, 95% CI 0.9-5.9, *P*_LMM_=0.067), a signal that remained apparent after controlling for *DQA1*\*01:03 (OR 2.5, 95% CI 1.0-6.1, *P*_LMM_=0.043). In the earlier report, the *DRB1\**14:01-*DQA1*\*01:01-*DQB1\**05:03 haplotype was found at higher frequency in cases of invasive GAS with severe systemic disease than either controls from the general population or cases of invasive GAS without severe systemic disease.^4^ While the former comparison is analogous to our analysis, the effect reported in that study was limited to the invasive GAS cases without NF, a finding that was not apparent from our data, with the caveat that the small numbers in both studies prevent a definitive conclusion. In our analysis, the signal at this haplotype is most likely explained by *DQB1*\*05:03, which was excluded from our primary analysis due to MAF 2.6% but showed the same borderline association with susceptibility (*P*_LMM_=0.064). Otherwise none of the previously implicated haplotypes were associated with susceptibility (Supplementary Table 4).

Limited effort has to date been documented investigating host genetic susceptibility to invasive GAS disease. As a starting point to further study in this area, we have demonstrated an association between the *HLA-DQA1*\*01:03 allele and susceptibility to invasive GAS disease in otherwise healthy children and adults. Importantly, we are encouraged by the high level of consistency of the *HLA-DQA1*\*01:03 association across a variety of sensitivity analyses including using data based on classical typing and use of an alternative control dataset. The presence of the rarer *HLA-DQA1*\*01:03 haplotype in one of 43 cases raises the possibility the association is driven by the allele rather than the background haplotype.

Beyond the earlier report linking the class II region to invasive GAS disease^4^, HLA has long been implicated in a range of infectious, autoimmune and other diseases.^12,13^. Moreover, the class II region has been a key finding in a number of recent GWAS of bacterial diseases including the somewhat analogous syndrome of invasive *Staphylococcus aureus* infection.^14,15^ The *DQA1*\*01:03 allele itself has not previously been implicated in susceptibility to infection but has been linked to several autoimmune and inflammatory diseases including primary sclerosis cholangitis^16^, systemic lupus erythematous^17^ and idiopathic achalasia.^18^ More recently, *DQA1*\*01:03 was part of one of several risk haplotypes that may potentially explain the HLA susceptibility locus in rheumatic heart disease, a post-infective complication of GAS infection.^19^ Thus, while further work will be required to fine-map the rheumatic heart disease association, it is possible that at least some genetic architecture may be shared across GAS diseases.

Interaction between HLA molecules and GAS superantigens has long been thought to play a key role in the pathogenesis of invasive GAS disease leading to activation of large numbers of T-cells.^3^ This process results in massive production of cytokines causing widespread tissue damage, disseminated intravascular thrombosis, and organ dysfunction which characterise the clinical picture.^20^ Crucially, binding of HLA by superantigens is largely antigen independent and usually occurs at residues outside the peptide-binding cleft.^21^ Moreover, SpeA, a key superantigen, binds with higher affinity to cell lines expressing *DQA1*\*01 alpha chains compared to *DQA1*\*03 or *DQA1*\*05 chains, to which very little binding was detected.^6^ By analogy to binding of staphylococcal enterotoxin B to DRB1^22^ and streptococcal superantigen to DQA1^23^, binding of SpeA to DQA1 is predicted to centre on a salt bridge formed between the glutamic acid at position 61 of SpeA and the lysine at position 42 of DQA1^24^, the latter widely termed position 39 in the superantigen literature in reference to the sequence of DRA.^22-24^ Tantalisingly, however, in *DQA1*\*01:03, the preceding arginine at position 41 is replaced by a second lysine, which could plausibly alter SpeA binding. Moreover, although heightened superantigen responsiveness might be expected to augment severity, it is also plausible that superantigens including SpeA may impair the acquisition of immunity to GAS thereby affecting susceptibility.^25-29^

Our study has three main limitations. First our sample size is small, especially by the standards of modern genetic research. Despite this, we propose that power is likely to be increased by our focus on patients with an extreme and well-defined phenotype of whom more than three quarters had NF, and by using a large number of controls, giving us an effective sample size of 167 in the primary analysis. Additionally, despite our more stringent upper age limit (65 vs 85 years), we include an equivalent number of previously healthy individuals with severe systemic disease (43 vs 44 cases) to that in the only comparable report in the literature.^4^ Thus it is of particular note that, although we have not made comparisons between severe and non-severe disease, we see very limited signal at the haplotypes reported to influence susceptibility in that report.^4^ One possible explanation for this difference is that, reflecting advances in genetic analysis since the publication of that report, our dataset underwent rigorous quality control including removal of individuals of outlying genetic ancestry limiting the risk of confounding due to issues such as differences in the genetic ancestry of cases and controls.^30^ Moreover, we analysed our data using linear mixed-models further curtailing confounding due to ancestry and relatedness^31^ which could plausibly contribute to the previously reported signals.^4^ This issue is also relevant to a recent study^29^ linking the *DQB1*\*06:02 allele to recurrent GAS-associated tonsillitis, the findings of which are difficult to interpret due inclusion of a mixture of Caucasian and Hispanic individuals without correction for ancestry at the analytical stage. Nonetheless, even allowing for the high-level of consistency across our sensitivity analyses, it is plausible that, owing to the small sample size, we may be over-estimating the effect of *DQA1*\*01:03 while being under-powered to detect other signals, including that linked to the rarer *DQB1*\*05:03 allele which might also influence susceptibility.

Second it is likely that having ascertained the cases through a patient group and a sample bank from a single institution they are not fully representative of invasive GAS disease in the general population, not least because those recruited through the patient group were all survivors who had predominantly suffered NF. That said, prospective recruitment at multiple institutions would be a costly and challenging endeavour which would have been hard to justify without the preliminary work presented here. Moreover, we consider the ascertainment of 34 otherwise healthy individuals with NF aged less than 65 years an accomplishment in itself, one that was possible only through the close involvement of a patient group.

Third with our current dataset we are unable to deconvolute whether the *HLA- DQA1*\*01:03 allele drives susceptibility to all invasive GAS disease or has a more specific effect on NF, although an effect on NF alone may be less likely given the weaker signal in the analysis limited to that subgroup. Similarly, due to limited data available on many cases, we are unable to ascertain whether the effect is dependent on variation in the bacteria, including the presence or absence of specific superantigen genes, or is influenced by other factors such as viral coinfections including influenza or varicella. Looking forward, however, we anticipate such questions will become answerable through large-scale prospective studies which will require collaborations involving investigators from multiple institutions and countries.

In summary, we have confirmed an association between class II polymorphism and invasive GAS disease, resolving it to a specific *DQA1* allele. Future research into the genetic basis of this devasting disease may bring about much-needed progress in development of vaccines or other therapeutics.

## Materials and Methods

Genetic data from cases of invasive GAS disease came from a newly genotyped sample collection, while genetic data from controls was from two existing datasets from earlier studies.

Cases aged less than 65 years without comorbidity were either survivors recruited retrospectively through the STREP GENE study (National Research Ethics Service Ref. 13/SC/0520) from a patient group called the Lee Spark NF Foundation (www.nfsuk.org.uk) or identified from a bank of samples at Imperial College London linked to limited clinical data that had been prospectively assembled from material surplus to diagnostic requirement (National Research Ethics Service Ref. 06/Q0406/20). Those recruited through the patient group had survived an episode of invasive GAS disease at a UK hospital since 1980 with microbiological confirmation obtained either through Public Health England or from the treating hospital. Participants submitted a saliva sample using Oragene® kits (DNA Genotek, Canada) from which DNA was extracted using the accompanying extraction kits. Those identified from the sample bank had been diagnosed with invasive GAS disease at the Imperial College Healthcare NHS Trust, London, UK, since 2006. DNA was extracted from stored tissue or serum using the Gentra® Puregene® Tissue kit (Qiagen®, USA) or QIAamp® Circulating Nucleic Acid kit (Qiagen). Cases were genome-wide genotyped using either the HumanCore platform (Illumina®, USA) or the Global Screening Array (Illumina). Controls for the primary analysis were children and adolescents recruited to various UK studies of vaccine efficacy for whom samples had been stored by the Oxford Vaccine Centre Biobank, University of Oxford, UK. These individuals had previously been genome-wide genotyped using the HumanOmniExpress platform (Illumina). Additional control data was available from the National Institute for Health Research Oxford Biobank including 5,544 individuals for whom validated HLA data were available.^9^

Quality control was undertaken using standard approaches^30^ but with an additional test^32^ aimed at identifying variants that differed between cases and controls due to the different platforms used for genotyping (Supplementary Table 1). During this process, seven cases were excluded, three on the basis of non-European ancestry, in addition to the six earlier exclude due to an age of 65 years or more (n=4) or co-morbidity (n=2). In total 119,134 variants genotyped on all three platforms were carried forward of which 434 were located in the extended major histocompatibility complex. For HLA imputation we used SNP2HLA software (version 1.0.3) without the default parameters using the prebuilt Type 1 Diabetes Genetics Consortium reference panel.^33^ With an overlap of 367 variants, we successfully imputed a total of 160 four-digit HLA alleles and 1,097 HLA amino-acid substitutions with imputation accuracy assessed exceeding 0.6 using the Beagle software^34^ (v3.0.4) R^2^ metric. Of these 47 four-digit alleles (27 in class II) and 869 amino acid substitutions (429 in class II) had minor allele frequency greater than 0.05. To minimise the effects of population structure and cryptic relatedness, we performed our primary analyses using linear mixed-models implemented in GCTA software^35^ (v1.26.0), limited to variants with MAF greater than 5% and with genotype represented by the dose of the minor allele estimated by imputation. We performed further analyses including estimation of effect sizes by transformation^36^ in R (v3.0) using amongst other tools the GenABEL^37^ and GMMAT packages^11^, and estimated the population attributable fraction as previously defined.^38^ To define three locus haplotypes in the class II region, we phased four-digit alleles using Phase^39^ software (v2.1.1) before extracting the probability of one or two copies of a given haplotype in each individual to define the dose of the minor allele. Finally, in a subset of samples, classical HLA typing of the class II locus using sequence-specific primer amplification was performed at the Transplant Immunology Laboratory at the Oxford Transplant Centre as previously described.^40^

## Acknowledgments

This research was supported by grants awarded to T.P. from the Medical Research Council (G1100449), the European Society of Clinical Microbiology & Infectious Diseases (Research Grant 2013) and the National Institute for Health Research (ACF-2016-20-001). J.C.K. is supported by a Wellcome Trust Investigator Award (204969/Z/16/Z). In addition, the genotyping undertaken by the Oxford Vaccine Group as part of EUCLIDS was funded through the European Union’s Seventh Framework Programme (EC-GA no. 279185) while the Oxford Biobank was funded through National Institute for Health Research Oxford Biomedical Research Centre, Oxford, UK. We thank the High-Throughput Genomics Group at the Wellcome Centre for Human Genetics for generating the genotyping data, subsidized by a core award from the Wellcome Trust (090532/Z/09/Z). We also acknowledge the support of the National Institute for Health Research Oxford Biomedical Research Centre, Oxford, UK, the National Institute for Health Research Imperial Biomedical Research Centre, London, UK and the National Institute for Health Research Health Protection Research Unit in Healthcare Associated Infections and Antimicrobial Resistance, Imperial College London, London, UK. None of these funders had any role in study design, data collection and analysis, decision to publish or preparation of the manuscript. Moreover, the views expressed here are those of the authors and not necessarily those of the National Health Service, the National Institute for Health Research or the Department of Health.

We thank the Lee Spark NF Foundation (www.nfsuk.org.uk; Charity No. 1088094) for assistance with recruitment as well as the patients and family members who took part in the STREP GENE study, as well as the volunteers who participated in the both the Oxford Vaccine Group (www.ovg.ox.ac.uk) and the Oxford Biobank (www.oxfordbiobank.org.uk) studies.

## Additional information

### Data availability

Genotype and phenotype data from invasive GAS cases underlying this manuscript have been deposited in the European Genome-phenome Archive (www.ega-archive.org) under accession number EGAS00001003421 with access permitted for further research on susceptibility to invasive GAS disease.

---

**Supplementary Table 1:** Quality Control of Genome-wide Genotyping Data

| **Process** | **Step** | **Threshold for exclusion** | **Removed** | **Carried Forward** |
| --- | --- | --- | --- | --- |
| Individual QC | Missingness | Missingness > 5% | 1 case<br>7 controls | 49 cases<br>2054 controls |
|  | Heterozygosity | Outliers removed<br>(see Supplementary Figure 3) | 2 cases<br>46 controls | 47 cases<br>2008 controls |
|  | Relatedness | Relatedness > 0.094 | 1 case<br>399 controls | 46 cases<br>1609 controls |
|  | Ancestry | Non-European outlier removed<br>(see Supplementary Figure 4) | 3 cases<br>69 controls | 43 cases<br>1540 controls |
| Variant QC | Missingness | Missingness > 0.01 | 73 variants | 119251 variants |
|  | Frequency | Minor allele frequency < 0.05 | 85 variants | 119166 variants |
| | Hardy-Weinberg | Hardy-Weinberg $P$-value < $10^{-10}$ | 9 variants | 119157 variants |
|  | Two Locus Test<sup>32</sup> | Outlier removed<br>(see Supplementary Figure 5) | 23 variants | 119134 variants |

**Supplementary Table 2:**
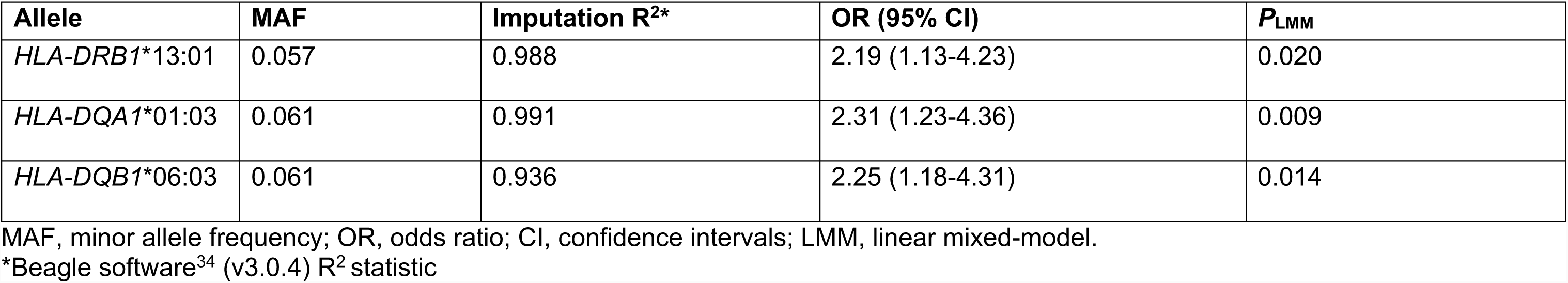
Association statistics for imputed HLA alleles associated with susceptibility

| **Allele** | **MAF** | **Imputation R<sup>2</sup>\*** | **OR (95% CI)** | **P<sub>LMM</sub>** |
| --- | --- | --- | --- | --- |
| *HLA-DRB1* *13:01 | 0.057 | 0.988 | 2.19 (1.13-4.23) | 0.020 |
| *HLA-DQA1* *01:03 | 0.061 | 0.991 | 2.31 (1.23-4.36) | 0.009 |
| *HLA-DQB1* *06:03 | 0.061 | 0.936 | 2.25 (1.18-4.31) | 0.014 |

MAF, minor allele frequency; OR, odds ratio; CI, confidence intervals; LMM, linear mixed-model.
\*Beagle software<sup>34</sup> (v3.0.4) R<sup>2</sup> statistic

**Supplementary Table 3:** Association statistics for imputed HLA amino acids associated with susceptibility

| Gene | IMGT Position | Amino acid | Direction | MAF | Imputation R<sup>2</sup>* | OR (95% CI) | P<sub>LMM</sub> |
| --- | --- | --- | --- | --- | --- | --- | --- |
| *HLA-A* | 9 | Tyrosine | Presence | 0.150 | 0.994 | 0.44<br>(0.20-0.96) | 0.041 |
| *HLA-C* | 35 | Glutamine | Presence | 0.136 | 0.999 | 1.86<br>(1.11-3.11) | 0.018 |
| *HLA-C* | 138 | Lysine | Presence | 0.136 | 0.997 | 1.85<br>(1.10-3.10) | 0.020 |
| *HLA-C* | 156 | Leucine-Arginine | Absence | 0.234 | 0.995 | 0.51<br>(0.27-0.94) | 0.031 |
| *HLA-C* | 275 | Glycine | Presence | 0.136 | 0.996 | 1.86<br>(1.11-3.11) | 0.018 |
| *HLA-DQA1* | 41 | Lysine | Presence | 0.061 | 0.990 | 2.32<br>(1.23-4.37) | 0.009 |
| *HLA-DQA1* | 130 | Alanine | Presence | 0.061 | 0.990 | 2.32<br>(1.23-4.36) | 0.009 |
| *HLA-DPB1* | 84 | Glycine | Absence | 0.315 | 0.976 | 1.55<br>(1.01-2.38) | 0.043 |
| *HLA-DPB1* | 205 | Valine | Absence | 0.124 | 0.766 | 2.01<br>(1.14-3.53) | 0.015 |
| *HLA-DPB1* | 215 | Isoleucine | Absence | 0.111 | 0.748 | 1.97<br>(1.09-3.55) | 0.024 |

IMGT, International Immunogenetics Information System; MAF, minor allele frequency; OR, odds ratio; CI, confidence intervals; LMM, linear mixed-model.
\*Beagle software<sup>34</sup> (v3.0.4) R<sup>2</sup> statistic

**Supplementary Table 4:** Association statistics for imputed previously implicated HLA haplotypes

| Haplotype | Direction* | MAF | Info† | OR (95% CI) | $P_{\text{LMM}}$ |
| --- | --- | --- | --- | --- | --- |
| *DRB1* *15:01- *DQA1* *01:02- *DQB1* *06:02 | Protective | 0.158 | 1.0 | 0.87 (0.47-1.64) | 0.67 |
| *DRB1* *03:01- *DQA1* *05:01- *DQB1* *02:01 | Protective | 0.142 | 1.0 | 0.97 (0.52-1.82) | 0.93 |
| *DRB1* *14:01- *DQA1* *01:01- *DQB1* *05:03 | Predisposition | 0.0262 | 1.0 | 2.36 (0.94-5.91) | 0.067 |
| *DRB1* *11:01- *DQA1* *05:01- *DQB1* *03:01 | Predisposition | 0.0753 | 1.0 | 0.79 (0.25-2.54) | 0.70 |
| *DRB1* *07:01- *DQA1* *03:01- *DQB1* *02:01 | Predisposition | 0 | - | - | - |

MAF, minor allele frequency; OR, odds ratio; CI, confidence intervals; LMM, linear mixed-model.
\*Reported in Kotb et al. 2002<sup>4</sup>
†SNPTEST software (v2.5.4) Info statistic

**Supplementary Figure 1:**
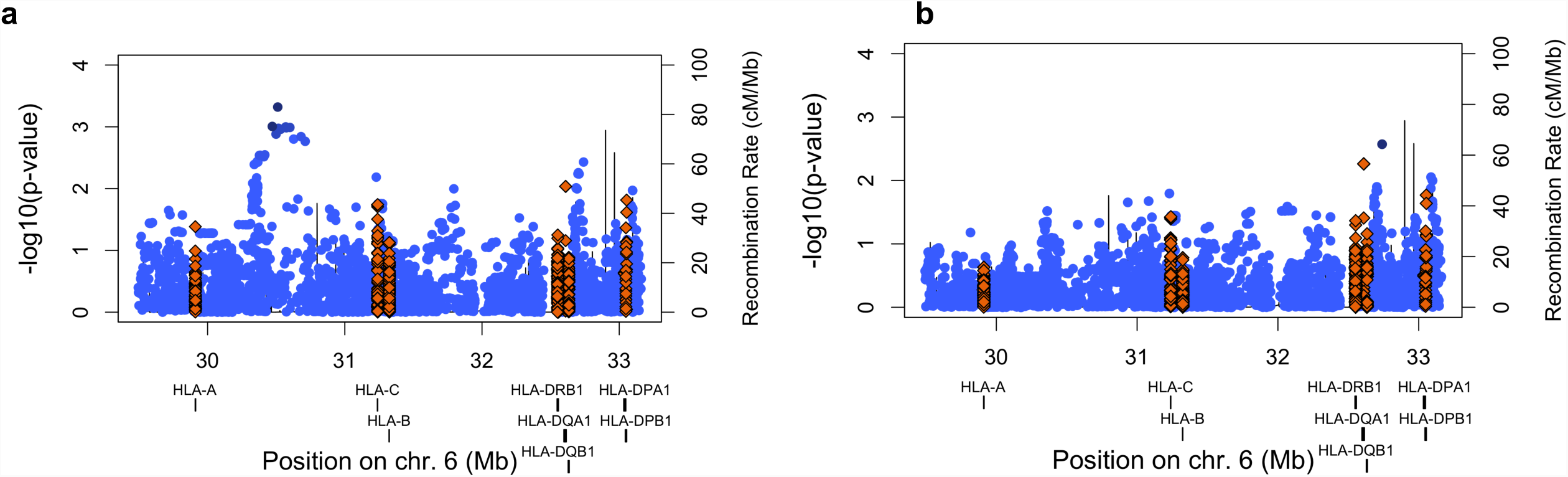
Association signals across the histocompatibility complex. Regional association plots are shown with genomic position plotted against the negative common logarithm of the p-value from a linear mixed-model analysis **(a)** before and **(b)** after conditioning on rs2524222. Amino acids are indicated by gold-coloured diamonds while SNPs are indicated blue circles with the depth of the blue proportional to the degree of linkage disequilibrium with the most associated variant. The recombination rate is shown as a line plotted on the right-hand y-axis.

**Supplementary Figure 2:**
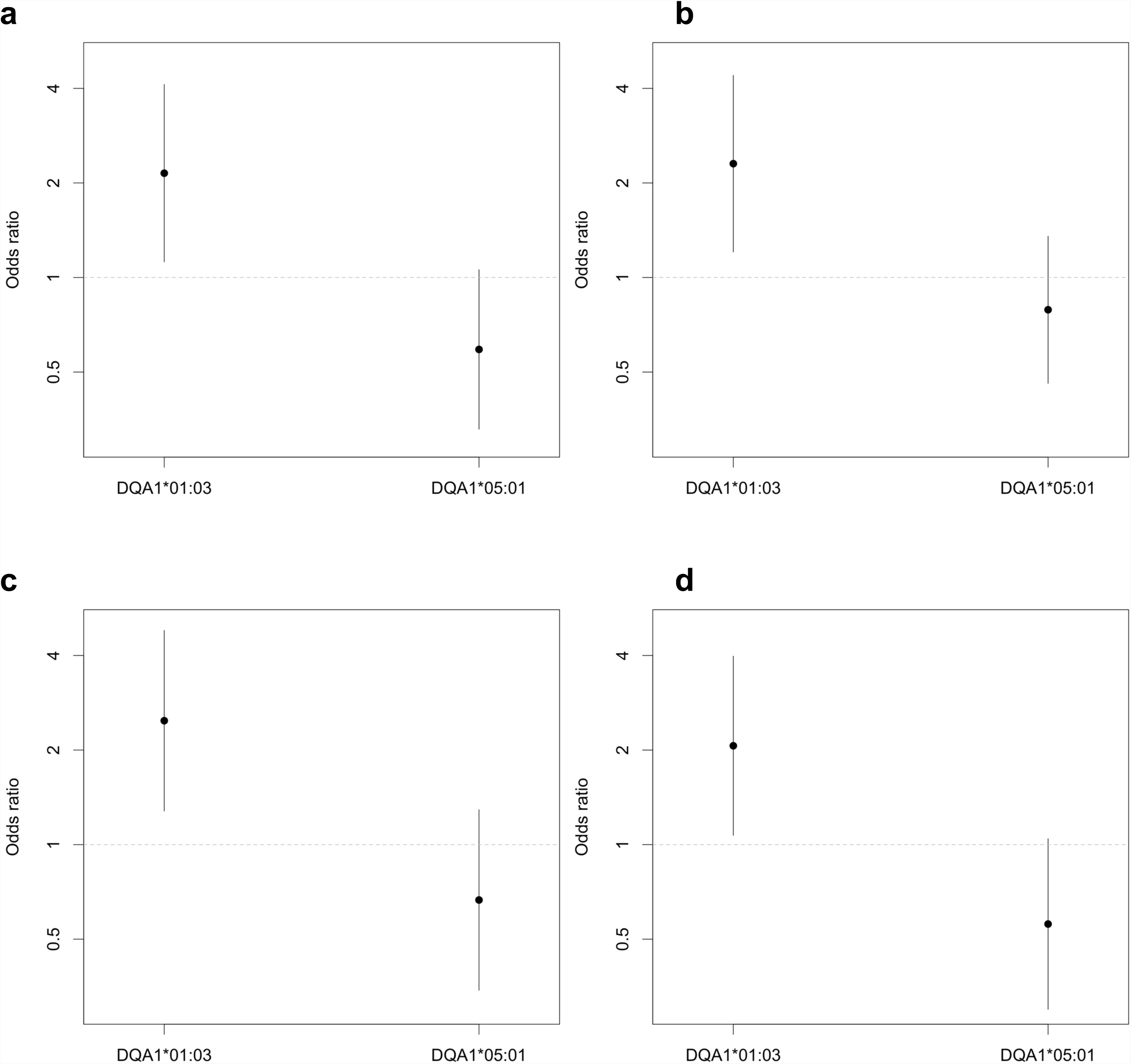
Effect size estimates for *HLA-DQA1*\*01:03 using alternative analytical approaches. Effect size estimates with confidence intervals are plotted for *HLA-DQA1*\*01:03 and *HLA-DQA1*\*05:01 based on a model including parameters for both alleles and sex using **(a)** a linear mixed-model with transformation^36^, **(b)** logistic regression with no additional parameters, **(c)** logistic regression with parameters for the first ten principal components^10^, and **(d)** a generalized linear mixed-model^11^.

**Supplementary Figure 3:**
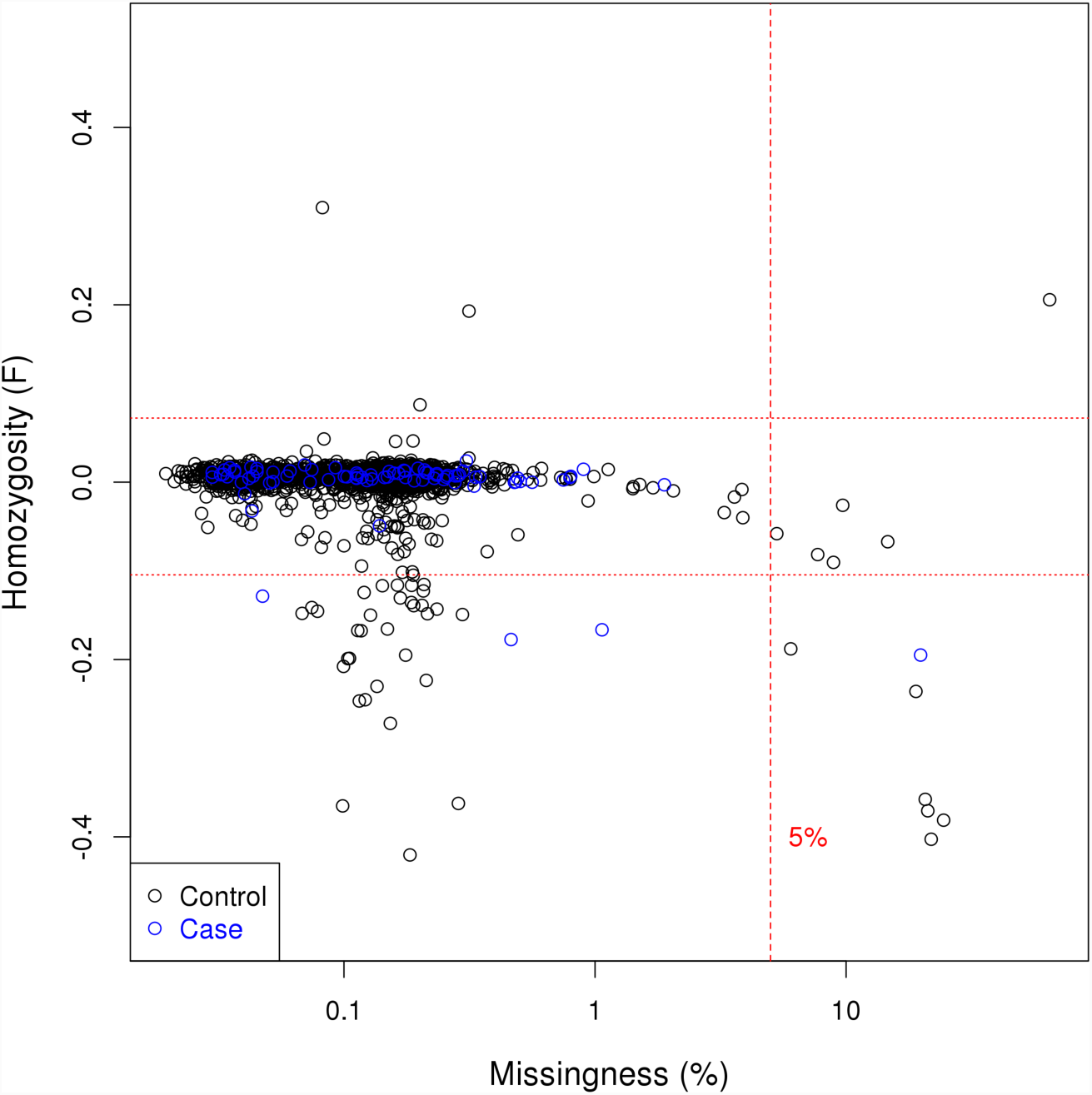
Assessment of heterogeneity and missingness in the study. Autosomal homozygosity is plotted against missingness on a logarithmic scale. Horizontal lines are drawn at two standard deviations above and three below the mean of autosomal homozygosity with missingness less than 5%.

**Supplementary Figure 4:**
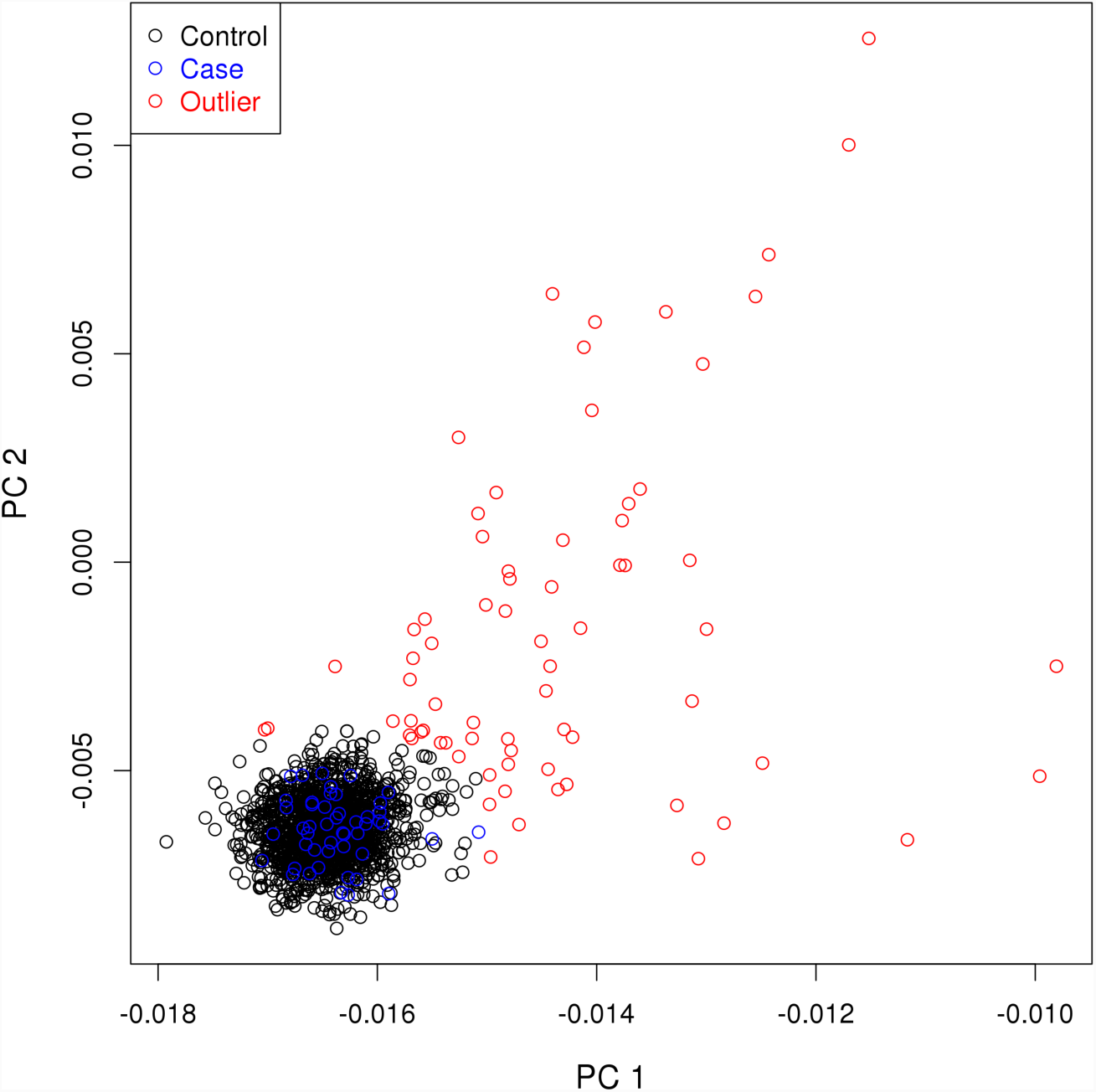
Principal component analysis of ancestry to definite European ancestry. Principal components analysis was run with HapMap consortium data^41^ from individuals of African and East Asian ancestry and outlying samples removed based on distance from the British European cluster.

**Supplementary Figure 5:**
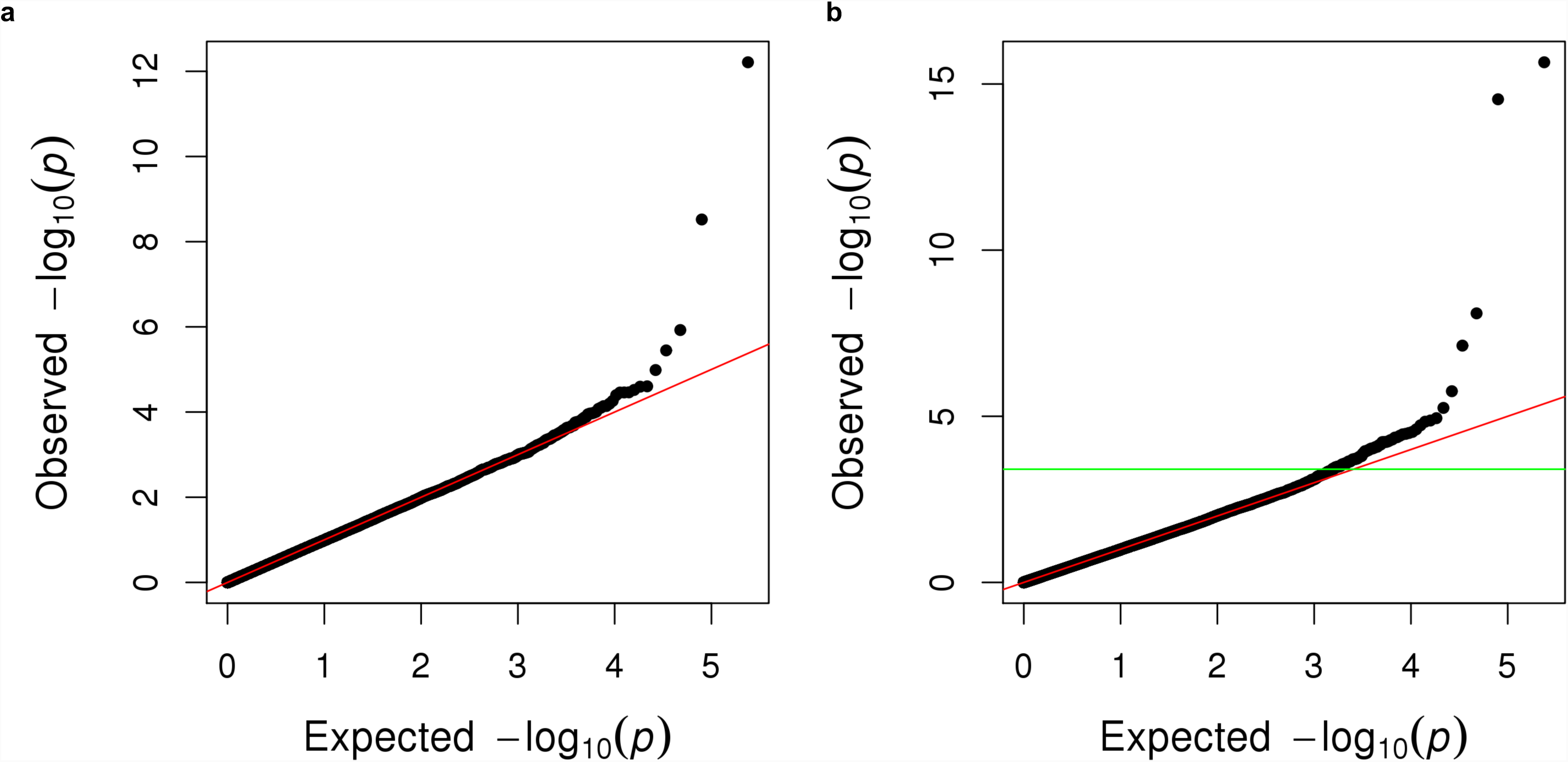
Removal of variants by the two locus quality control test. Quantile-quantile plots are shown for genome-wide susceptibility analysis performed with **(a)** unadjusted linear regression and **(b)** the genome-wide linear model-based quality control test. In the latter, the horizontal green line is drawn at the point at which the negative common logarithm of the observed *P*-value exceeds the expected value by greater than 0.2 (based on Lee *et al.*^32^). Variants were excluded when the test statistic calculated in both directions exceeded this threshold.

## References

1. Lamagni TL, Darenberg J, Luca-Harari B, Siljander T, Efstratiou A, Henriques-Normark B et al. Epidemiology of severe Streptococcus pyogenes disease in Europe. J Clin Microbiol 2008; 46: 2359–2367.

2. Lamagni TL, Neal S, Keshishian C, Powell D, Potz N, Pebody R et al. Predictors of death after severe Streptococcus pyogenes infection. Emerging Infect Dis 2009; 15: 1304–1307.

3. Musser JM, Shelburne SA. A decade of molecular pathogenomic analysis of group A Streptococcus. J Clin Invest 2009; 119: 2455–2463.

4. Kotb M, Norrby-Teglund A, McGeer A, El-Sherbini H, Dorak MT, Khurshid A et al. An immunogenetic and molecular basis for differences in outcomes of invasive group A streptococcal infections. Nat Med 2002; 8: 1398–1404.

5. Imanishi K, Igarashi H, Uchiyama T. Relative abilities of distinct isotypes of human major histocompatibility complex class II molecules to bind streptococcal pyrogenic exotoxin types A and B. Infect Immun 1992; 60: 5025–5029.

6. Llewelyn M, Sriskandan S, Peakman M, Ambrozak DR, Douek DC, Kwok WW et al. HLA class II polymorphisms determine responses to bacterial superantigens. J Immunol 2004; 172: 1719–1726.

7. Sriskandan S, Unnikrishnan M, Krausz T, Dewchand H, Van Noorden S, Cohen J et al. Enhanced susceptibility to superantigen-associated streptococcal sepsis in human leukocyte antigen-DQ transgenic mice. J Infect Dis 2001; 184: 166–173.

8. Kasper KJ, Zeppa JJ, Wakabayashi AT, Xu SX, Mazzuca DM, Welch I et al. Bacterial Superantigens Promote Acute Nasopharyngeal Infection by Streptococcus pyogenes in a Human MHC Class II-Dependent Manner. PLoS Pathog 2014; 10: e1004155.

9. Neville MJ, Lee W, Humburg P, Wong D, Barnardo M, Karpe F et al. High resolution HLA haplotyping by imputation for a British population bioresource. Hum Immunol 2017; 78: 242–251.

10. Price AL, Patterson NJ, Plenge RM, Weinblatt ME, Shadick NA, Reich D. Principal components analysis corrects for stratification in genome-wide association studies. Nat Genet 2006; 38: 904–909.

11. Chen H, Wang C, Conomos MP, Stilp AM, Li Z, Sofer T et al. Control for Population Structure and Relatedness for Binary Traits in Genetic Association Studies via Logistic Mixed Models. Am J Hum Genet 2016; 98: 653–666.

12. Chapman SJ, Hill AVS. Human genetic susceptibility to infectious disease. Nat Rev Genet 2012; 13: 175–188.

13. Trowsdale J, Knight JC. Major histocompatibility complex genomics and human disease. Annu Rev Genomics Hum Genet 2013; 14: 301–323.

14. DeLorenze GN, Nelson CL, Scott WK, Allen AS, Ray GT, Tsai A-L et al. Polymorphisms in HLA Class II Genes Are Associated With Susceptibility to Staphylococcus aureus Infection in a White Population. J Infect Dis 2016; 213: 816–823.

15. Cyr DD, Allen AS, Du GJ, Ruffin F, Adams C, Thaden JT et al. Evaluating genetic susceptibility to Staphylococcus aureus bacteremia in African Americans using admixture mapping. Genes Immun 2017; 18: 95–99.

16. Liu JZ, Hov JR, Folseraas T, Ellinghaus E, Rushbrook SM, Doncheva NT et al. Dense genotyping of immune-related disease regions identifies nine new risk loci for primary sclerosing cholangitis. Nat Genet 2013; 45: 670–675.

17. Langefeld CD, Ainsworth HC, Cunninghame Graham DS, Kelly JA, Comeau ME, Marion MC et al. Transancestral mapping and genetic load in systemic lupus erythematosus. Nat Commun 2017; 8: 16021.

18. Gockel I, Becker J, Wouters MM, Niebisch S, Gockel HR, Hess T et al. Common variants in the HLA-DQ region confer susceptibility to idiopathic achalasia. Nat Genet 2014; 46: 901–904.

19. Gray L-A, D’Antoine HA, Tong SYC, McKinnon M, Bessarab D, Brown N et al. Genome-Wide Analysis of Genetic Risk Factors for Rheumatic Heart Disease in Aboriginal Australians Provides Support for Pathogenic Molecular Mimicry. J Infect Dis 2017; 216: 1460–1470.

20. Walker MJ, Barnett TC, McArthur JD, Cole JN, Gillen CM, Henningham A et al. Disease manifestations and pathogenic mechanisms of group a streptococcus. Clin Microbiol Rev 2014; 27: 264–301.

21. Sriskandan S, Altmann DM. The immunology of sepsis. J Pathol 2008; 214: 211–223.

22. Jardetzky TS, Brown JH, Gorga JC, Stern LJ, Urban RG, Chi YI et al. Threedimensional structure of a human class II histocompatibility molecule complexed with superantigen. Nature 1994; 368: 711–718.

23. Sundberg E, Jardetzky TS. Structural basis for HLA-DQ binding by the streptococcal superantigen SSA. Nat Struct Biol 1999; 6: 123–129.

24. Papageorgiou AC, Collins CM, Gutman DM, Kline JB, O’Brien SM, Tranter HS et al. Structural basis for the recognition of superantigen streptococcal pyrogenic exotoxin A (SpeA1) by MHC class II molecules and T-cell receptors. EMBO J 1999; 18: 9–21.

25. Eriksson BK, Andersson J, Holm SE, Norgren M. Invasive group A streptococcal infections: T1M1 isolates expressing pyrogenic exotoxins A and B in combination with selective lack of toxin-neutralizing antibodies are associated with increased risk of streptococcal toxic shock syndrome. J Infect Dis 1999; 3 180: 410–418.

26. Basma H, Norrby-Teglund A, Guédez Y, McGeer A, Low DE, El-Ahmedy O et al. Risk factors in the pathogenesis of invasive group A streptococcal infections: role of protective humoral immunity. Infect Immun 1999; 67: 1871–1877.

27. Proft T, Fraser JD. Streptococcal Superantigens: Biological properties and potential role in disease. In: Ferretti JJ, Stevens DL, Fischetti VA, eds. Streptococcus pyogenes : Basic Biology to Clinical Manifestations. Oklahoma City: University of Oklahoma Health Sciences Center, 2016.

28. Davies FJ, Olme C, Lynskey NN, Turner CE, Sriskandan S. Streptococcal superantigen-induced expansion of human tonsil T cells leads to altered T follicular helper cell phenotype, B cell death, and reduced immunoglobulin release. biorXiv 2018. doi:10.1101/395608.

29. Dan JM, Havenar-Daughton C, Kendric K, Al-Kolla R, Kaushik K, Rosales SL et al. Recurrent group A Streptococcus tonsillitis is an immunosusceptibility disease involving antibody deficiency and aberrant TFH cells. Sci Transl Med 2019; 11: eaau3776.

30. Anderson CA, Pettersson FH, Clarke GM, Cardon LR, Morris AP, Zondervan KT. Data quality control in genetic case-control association studies. Nat Protoc 2010; 5: 1564–1573.

31. Yang J, Zaitlen NA, Goddard ME, Visscher PM, Price AL. Advantages and pitfalls in the application of mixed-model association methods. Nat Genet 2014; 46: 100–106.

32. Lee SH, Nyholt DR, Macgregor S, Henders AK, Zondervan KT, Montgomery GW et al. A Simple and Fast Two-Locus Quality Control Test to Detect False Positives Due to Batch Effects in Genome-Wide Association Studies. Genet Epidemiol 2010; 34: 854–862.

33. Jia X, Han B, Onengut-Gumuscu S, Chen W-M, Concannon PJ, Rich SS et al. Imputing Amino Acid Polymorphisms in Human Leukocyte Antigens. PLoS ONE 2013; 8: e64683.

34. Browning BL, Browning SR. A Unified Approach to Genotype Imputation and Haplotype-Phase Inference for Large Data Sets of Trios and Unrelated Individuals. The American Journal of Human Genetics 2009; 84: 210–223.

35. Yang J, Lee SH, Goddard ME, Visscher PM. GCTA: A tool for genome-wide complex trait analysis. Am J Hum Genet 2011; 88: 76–82.

36. Lloyd-Jones LR, Robinson MR, Yang J, Visscher PM. Transformation of Summary Statistics from Linear Mixed Model Association on All-or-None Traits to Odds Ratio. Genetics 2018; 208: 1397–1408.

37. Aulchenko YS, Ripke S, Isaacs A, van Duijn CM. GenABEL: an R library for genome-wide association analysis. Bioinformatics 2007; 23: 1294–1296.

38. Witte JS, Visscher PM, Wray NR. The contribution of genetic variants to disease depends on the ruler. Nat Rev Genet 2014; 15: 765–776.

39. Stephens M, Smith NJ, Donnelly P. A new statistical method for haplotype reconstruction from population data. Am J Hum Genet 2001; 68: 978–989.

40. Welsh K, Bunce M. Molecular typing for the MHC with PCR-SSP. Rev Immunogenet 1999; 1: 157–176.

41. The International HapMap Consortium. A haplotype map of the human genome. Nature 2005; 437: 1299–320.

